# Co-infection with *Leptomonas seymouri* enhances macrophage survival and promotes intracellular parasite persistence during *Leishmania donovani* infection

**DOI:** 10.64898/2026.02.23.707439

**Authors:** Sayantan Das, Pritam Dey Sarkar, Subhajit Biswas

## Abstract

**Background:** *Leishmania donovani* (LD) is an obligate intracellular parasite that survives and replicates within macrophages. *Leptomonas seymouri* (LS), a traditionally monoxenous trypanosomatid, has been repeatedly co-isolated with LD from kala-azar cases in India, often together with *Leptomonas seymouri* narna-like virus 1 (Lepsey NLV1). Whether LS can survive and replicate within mammalian macrophages, and how co-infection influences parasite and virus dynamics, remain unresolved.

**Methods and Findings:** Intracellular survival, *replication*, and revival of LS alone and during co-infection with LD were systematically evaluated using murine (RAW 264.7) and human (THP-1) macrophages. Quantitative ITS1 PCR demonstrated significant increases in intracellular parasite DNA over 48-168 hours post-infection (h.p.i), indicating active replication rather than persistence. Reduced extracellular count suggested restricted cell lysis and enhanced macrophage survival in co-infection compared to mono-infections. Giemsa staining confirmed intracellular localization of LS. Amastigotes from infected macrophages revived as motile promastigotes upon transformation, whereas extracellular parasites failed to revive beyond 48 hours, confirming macrophages as the exclusive niche for prolonged viability. Co-infection dampened macrophage IL-12 (p40) production, augmenting macrophage survival. MTT assay confirmed the same. In THP-1, co-infection resulted in a marked increase in intracellular LS count and viral load than LS mono-infection at 96 and 168 h.p.i. relative to 48 h.p.i., suggesting that LD co-infection favors higher intracellular virus titres.

**Conclusions:** Our *findings* demonstrated that LS could replicate within mammalian macrophages and persist during mono-or LD co-infection. The identification of a stable LD-LS-virus interaction highlights a previously underappreciated “triple-pathogen” biology with potential implications for LD pathogenesis.

## 1. Introduction

*Leishmania* parasites are obligate intracellular pathogens that rely on macrophages for their survival, replication, and differentiation **(Liu and Uzonna, 2012).** Despite the extreme challenges posed by the intracellular environment, *Leishmania* are among the few protozoa capable of surviving and multiplying under such harsh conditions **(Gregory and Olivier, 2005)**. *Leptomonas seymouri* (*L. seymouri*; LS), typically a monoxenous trypanosomatid that infects insects, is generally not capable of infecting humans **(Thakur et al., 2020).** However, several reports from India indicated that this parasite could co-infect humans alongside *Leishmania donovani* (*L. donovani*; LD) in cases of visceral leishmaniasis (VL), cutaneous leishmaniasis and post-kala-azar dermal leishmaniasis (PKDL) **(Ghosh et al., 2012a; Selvapandiyan et al., 2015; Thakur et al., 2020; Dixit et al., 2021).**

An in-depth study had been previously conducted to evaluate the infective potential of LS. It was reported that despite several genetic adaptations, LS remained primarily a monoxenous organism and couldn’t infect mammalian macrophages such as J774 or BMM , either on its own or alongside LD **(Kraeva et al., 2015).** Others implicated that it might infect mammals under specific conditions, for instance, when the host’s immune system was compromised by another pathogen such as LD or HIV **(Selvapandiyan et al., 2015).** Interestingly, a study published in 1981 examined the interactions between LD and *Leptomonas costoris* within peritoneal macrophages from hamster. It was observed that LD differentiated into amastigotes and survived within cultured macrophages, whereas *L. costoris* also differentiated into amastigotes but failed to survive. Additionally, macrophage cultures infected with the infectious strain of LD retained 50% more *L. costoris* compared to control macrophages that were infected solely with *L. costoris* **(Kutish and Janovy, 1981).**

Macrophages serve as both the primary niche for *Leishmania* survival and replication, and as a key effector population in parasite clearance **(DO, 1992)**. However, the cellular consequences of LS infection on macrophages remained largely unexplored, highlighting a critical knowledge gap that warranted systematic investigation.

The presence of *Leishmania* RNA virus (LRV) is known to enhance the persistence and worsen the pathogenesis of *Leishmania sp*.**(Adaui et al., 2016; Rossi et al., 2017).** Unexpectedly, LS was found to harbour a distinct protozoan virus-*Leptomonas seymouri* narna-like virus 1 (Lepsey NLV1), an RNA virus of approximately 4.5 kb **(Lye et al., 2016)**. A study from India reported that >90% of LD laboratory isolates from clinical samples of VL were co-infected with LS. Lepsey NLV1 was predominantly present in the LS-positive samples (75% of the laboratory isolates) **(Sukla et al., 2017).** The same laboratory further reported that LS and Lepsey NLV1 could be even detected alongside LD in archived serum samples of kala-azar or PKDL patients. It also suggested that Lepsey NLV1 might enhance the persistence of LD and dampen immune responses by down-regulating IL-18 **(Sukla et al., 2022)** ,which also corroborated with the results from a subsequently published LRV2+ *L. major* study **(Mirabedini et al., 2023).** Leishmania RNA viruses (LRVs) had been known to exacerbate *Leishmania* pathogenesis. In comparison, whether or not Lepsey NLV1 is linked to pathogenesis or persistence of VL /PKDL (“triple pathogen phenomenon”) warrants further investigation.

In this study, we aimed to systematically investigate the infective potential of LS in mammalian macrophages and to determine whether LS can survive, replicate, and persist intracellularly, either alone or in co-infection with LD. We further sought to evaluate the impact of LD-LS co-infection on parasite population dynamics, host macrophage responses, and the persistence of Lepsey NLV1 virus. Through a combination of molecular, cellular, and functional assays, this study attempts to address the unresolved question of whether LS can adapt to a mammalian intracellular niche and contribute to a potential “triple-pathogen” interaction relevant to disease pathogenesis.

## 2. MATERIALS AND METHODS

### 2.1. Parasite culture

*Leishmania donovani* (MHOM/IN/1983/AG83) and *Leptomonas seymouri* (ATCC) promastigotes were cultured at 22°C in M199 (Sigma) with 10% heat-inactivated fetal bovine serum (FBS) (Gibco) and 1% L-glutamine-penicillin-streptomycin solution (Sigma). Promastigotes were harvested on the 6^th^ or 7^th^ day of the stationary phase of growth for *in vitro* infection. Laboratory isolates of LD used in this study was confirmed Lepsey NLV1 negative by virus specific PCR.

### 2.2. Murine macrophage culture

Murine macrophage cell line RAW 264.7 was cultured at 37°C with 5% CO_2_ in DMEM (Dulbecco’s modified Eagle’s medium) (Sigma) supplemented with 10% heat-inactivated fetal bovine serum (FBS) (DMEM-10) (Gibco) and 1% L glutamine-penicillin-streptomycin solution (Sigma).

For infection experiments, cells were plated in 12-well flat bottom plates (Nunclon Delta Surface^TM^, Thermo Scientific) with 2x10^5^ cells /well and incubated at 37°C with 5% CO_2_ for 48 hours (h). Cells were counted at 8x10^5^ /well at 80% confluency.

### 2.3. Human monocyte culture

Human monocyte cell line THP-1 was cultured at 37°C with 5% CO_2_ in RPMI-1640 medium (Sigma) supplemented with 10% heat-inactivated fetal bovine serum (FBS) (Gibco) and 1% L glutamine-penicillin-streptomycin solution (Sigma).

### 2.4. THP-1 activation

For infection experiments, cells were plated in 6-well flat bottom plates (Nunclon Delta Surface^TM^, Thermo Scientific) with 5x10^5^ cells /well. Before infection, cells were stimulated with 100 nM /well of 4α-phorbol 12-myristate 13-acetate (PMA) (Sigma, USA) to induce differentiation into macrophage-like cells. The cultures were then incubated for 24 h at 37°C in a humidified atmosphere containing 5% CO_2_. Following differentiation, wells were gently washed twice with culture medium (without PMA) to remove any non-adherent cells.

### 2.5. *In vitro* infection

Accordingly, macrophages were infected with 10^7^ stationary phase promastigotes of LD, LS and 2:1 or 10:1 co-culture (LD: LS) at a multiplicity of infection (MOI) of 10:1 (parasite: Mφ ratio) and incubated at 37°C with 5% CO_2_ for 4 h. Cells were then washed twice with media to remove non-adherent parasites and incubated with media for 48 h to 168 h. At the end of post-infection incubation period, the cells were processed for further analysis.

### 2.6. Extraction of purified Lepsey NLV1 and *in vitro* infection with virus

For extraction of purified Lepsey NLV1 virus, approx. 10^8^ LS parasites were frozen at - 80°C for 3 h, followed by thawing at 37°C. Sonication was performed using three pulses of 10 seconds (s) each. The samples were then centrifuged at 3000 rpm for 10 minutes (min) to pellet down the ruptured LS parasites. The resulting supernatant, containing Lepsey NLV1, was collected, passed through a 0.22 μm syringe filter to remove residual cellular debris, and stored at -80°C, in 500 μL aliquots for downstream applications.

For *in vitro* infection, RAW 264.7 macrophage cells were infected with purified Lepsey NLV1 at an MOI of 10:1 and incubated at 37 °C with 5% CO_2_ for 4 h. The inoculum was not removed after virus infection and supplemented with DMEM-1 for further incubation. After 48-, 72-and 96-hours post-infection (h.p.i.), cell supernatant and lysate were collected for further assays.

### 2.7. Macrophage infected parasite revival assay

To check the parasite’s viability and if they can revive and transform from infected macrophages, 10^6^ RAW 264.7 macrophages were infected with only LD, LS or a 2:1 or 10:1 (LD: LS) co-culture at an MOI of 10:1. After 4 h of infection, non-adhered parasites were removed by washing the cells twice with 1X PBS. Cells were maintained in DMEM-10 at 37°C with 5% CO_2_ for 48 to 168 h. After every incubation period, the supernatant was collected. Cells were scrapped in 500 µL of DMEM-10 and were added to 5 mL Schneider’s Drosophila medium (Gibco) with 20% FBS. Flasks were incubated at 22°C and incubated for at least 48 h to check the transformation of parasites from infected macrophages.

To exclude any possibility of parasite viability in DMEM-10, 10^7^ parasites /well were maintained in DMEM-10 at 37°C for 48-168 h. Subsequently, 2 x 10^6^ parasites /time point were transferred to Schneider’s Drosophila medium at 22°C to assess revival capacity at different time points. Parasite load was quantified 9 days post inoculation in Schneider’s Drosophila media using ITS1 qPCR. In another set of experiment, 10^6^ viable LS parasites /well were inoculated into 12-well plates and incubated at 37°C in DMEM-10 for 8 days. For further verification, 10^6^ LD and 10^6^ LS parasites were cultured in both DMEM-10 and RPMI-10 at 37°C for a period of 24, 72-and 168 h time points. Both microscopic and molecular analysis (ITS1 PCR) were performed to check the replication of parasites. Parasite load was determined from densitometric analysis of specific bands on gel using ImageJ software.

### 2.8. DNA extraction and ITS1 gene PCR

DNA was extracted from 200 µL of parasite infected cell supernatant or cell lysate using a QIAamp DNA Mini kit (Qiagen). The quality and concentration of DNA was assessed using a NanoDrop spectrophotometer (Thermo Scientific, USA).

An rDNA-ITS1 fragment was directly amplified from the isolated DNA using ITS1-FP (5’-CTGGATCATTTTCCGATG-3’) and ITS1-RP (5’-TGATACCACTTATCGCACTT-3’) primers **(El Tai et al., 2000)**. The PCR protocol included an initial denaturation at 94°C for 10 min, followed by 40 cycles consisting of denaturation at 94°C for 1 min, annealing at 54°C for 1 min, and extension at 72°C for 1 min. A final elongation step was performed at 72°C for 10 min. PCR products were analysed using 1% agarose gel containing SYBRsafe^TM^ (Invitrogen) for nucleic acid staining. DNA from the LD and LS was used as a positive control, while nuclease-free water served as the non-template control (NTC). The ITS1 PCR assay generated species-specific PCR products of distinct sizes, yielding a 320 bp product for LD and a 418 bp product for LS. In co-infected samples, simultaneous detection of both PCR products enabled differentiation and confirmation of the presence of both parasite species within the same sample.

### 2.9. Real Time PCR and Image J Analysis

A quantitative real-time PCR assay was designed to quantify the parasite DNA copy number in the samples using a SYBR Green dye-based quantitative qPCR with the Luna Universal PCR reagent (New England Biolabs). Equal amounts of DNA from each experimental condition were used for qPCR analysis. A 320 bp or 418 bp fragment of the ITS 1 gene was amplified using the same primers as mentioned above. The reactions were carried out on a StepOne^TM^ system (Applied Biosystems, Thermo).

PCR products specificity was confirmed by resolving qPCR products on 1% agarose gel, which consistently yielded bands of the expected sizes. To quantify species composition in co-infected samples, band intensities corresponding to LD and LS (including serially-diluted standards and experimental samples) were extracted from gel images and subjected to densitometric analysis using ImageJ. The LD: LS ratios derived from this analysis were subsequently used to estimate the relative contribution of each parasite to the total burden under co-infection conditions.

### 2.10. MTT Cell Viability Assay

Murine macrophages were seeded in 96-well plates at 1 × 10^4^ cells /well and allowed to adhere overnight. Cells were washed with 1X PBS and infected with LD, LS or co-infection mixtures (LD: LS ratios of 2:1, 5:1, and 10:1) at an MOI of 10:1. After 4 h, non-internalized parasites were removed by 1X PBS washing, and cells were incubated in fresh complete DMEM-10 for 96 h at 37°C with 5% CO_2_.

Cell viability was determined using the MTT assay. Post infection, cells were washed with 1X PBS and medium was replaced with 90 µL phenol red-free DMEM, followed by addition of 10 µL MTT (5 mg /mL), and incubation for 4 h at 37°C in the dark. Formazan crystals were dissolved in 100 µL DMSO, and absorbance was measured at 595 nm.

Cell viability (%) was calculated relative to uninfected controls using the formula: [(Abs_sample − Abs_blank) /(Abs_control − Abs_blank)] × 100; Abs = Absorbance. Wells without cells served as blanks while uninfected cells served as positive controls.

### 2.11. RNA Extraction and cDNA synthesis

Total RNA was extracted from infected macrophage cell supernatant or lysate using standard TRIzol method following manufacturer’s protocol (PureLink RNA Minikit, Invitrogen). The concentration and purity of RNA was determined using a Nanodrop One spectrophotometer (Thermo Scientific, USA).

The extracted RNA was subjected to cDNA synthesis and virus specific nested PCR using primers and protocol as described previously **(Sukla et al., 2022)**. PCR products were separated using 1% agarose gel electrophoresis.

### 2.12. Real Time PCR

The virus RNA copy number in the nucleic acid isolated from the samples were ascertained using qRT-PCR. The primers CT2-F and CT2-R were used to amplify a 338 bp segment of the Lepsey NLV1 gene using protocol as described elsewhere **(Sukla et al., 2022)**. Specific amplified products were visualized in 1% agarose gel electrophoresis with SYBRsafe^TM^ for nucleic acid staining.

### 2.13. Giemsa Staining

Cells were plated in a 4-well chamber (Genetix Biotech) tissue culture slide with 5x10^4^ cells (RAW 264.7) or 10^5^ (THP-1) and incubated at 37°C with 5% CO_2_ overnight. Macrophages were infected with the stationary phase of LD, LS or 10:1 co-culture (LD: LS) promastigotes at an MOI of 10:1 and incubated at 37°C with 5% CO_2_ for 4 h. Cells were then washed twice with media to remove non-adherent parasites and incubated with media for 72 h. After the incubation period, cells were fixed with methanol and stained with Giemsa. The stained slides were visualized under an optical microscope (EVOS microscope) for determining the infection of macrophages at 1000X magnification.

### 2.14. IL-12 (p40) ELISA

Quantitative IL-12 (p40) ELISA (BD OptEIA™, Cat. No.-555165) was performed as per manufacturer’s protocol to determine cytokine levels from cell supernatant of infected RAW 264.7 cells at 72 h.p.i. Standard curve was prepared using dilutions of the respective recombinant cytokine provided in the kit.

### 2.15. Statistical Analysis

Statistical analysis was performed using GraphPad Prism version 8.0 (GraphPad Software, USA). Data are expressed as mean ± SD unless otherwise indicated. Statistical tests were selected based on experimental design and data distribution. Differences between two groups were evaluated using unpaired two-tailed Student’s t-test. For comparisons involving more than two groups, one-way ANOVA followed by Dunnett or Tukey’s post hoc tests were used. Each experiment was performed using at least three independent biological replicates. A p value of < 0.05 was considered statistically significant.

## 3. RESULTS

### 3.1. Monoxenous parasite *Leptomonas seymouri* harbouring Lepsey NLV1 could efficiently infect and replicate in mammalian macrophage cells

#### 3.1.1. LD-LS co-infection resulted in higher intra-cellular parasite load but better macrophage survival compared to mono-infections in RAW 264.7 cells

ITS1 PCR based quantification revealed that the intracellular LS parasite load within RAW 264.7 cells increased over time from 48-to 96-and 168 h.p.i. by ∼ 3 and 5.3-fold respectively. LS load significantly increased at 168 h.p.i compared to 48 h.p.i. The ∼2-fold rise in IC LS DNA at 168 h.p.i. compared to the initial inoculum (10^7^ LS; though two-times washed after 4 h of cellular internalization) confirmed true intracellular replication rather than residual carry-over. In case of co-infection, the total intracellular parasite load also significantly increased over time at 96-and 168 h.p.i compared to 48 h.p.i by ∼4.9 and ∼7.3-fold respectively. Intracellular LD parasite load also significantly increased over 96-and 168 h.p.i compared to 48 h.p.i but to lower counts compared to LS mono-or co-infection. (Fig. 1A) and (Supplementary Fig. S1).

**Fig. 1.**
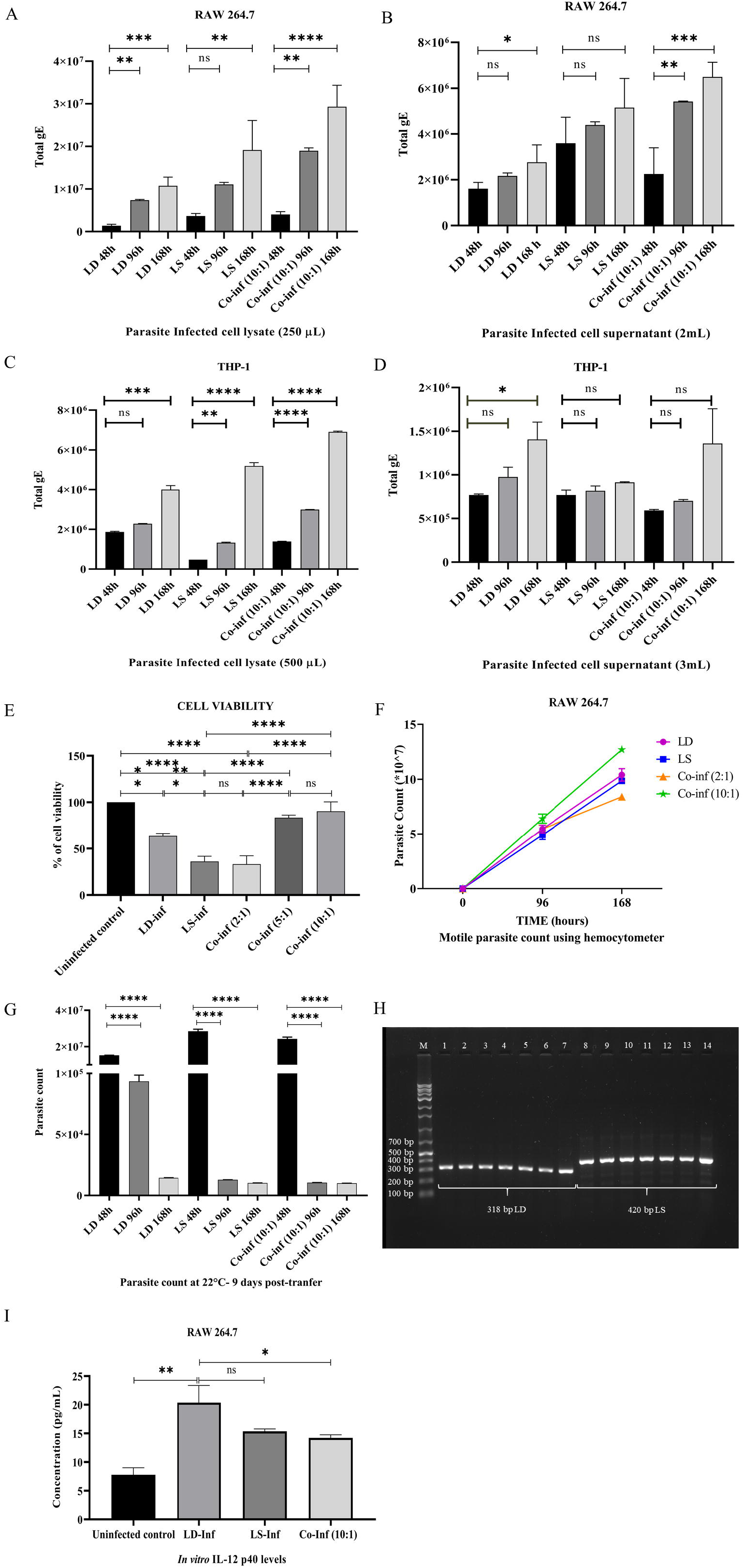
Quantification of parasite load from infected macrophage cells. Parasite load was assessed in culture **(A)** cell lysate and **(B)** supernatant from parasite infected RAW 264.7 cells at 48-, 96-and 168 hours post-infection (h.p.i.). Parasite load quantified from infected THP-1 macrophage **(C)** cell lysate and **(D)** supernatant at 48-,96-and 168-h.p.i. **(E)** Cell viability of murine macrophages infected with LD, LS, and LD–LS co-infections at varying ratios, assessed using MTT assay. **(F)** Motile parasite counts from transformation /subculture flasks at 12 days post-transfer, derived from infected macrophage harvests collected at 0-, 96-and 168 h.p.i and monitored using a hemocytometer. **(G)** Approximately 10 parasites /well were maintained in DMEM-10 at 37°C for 48-, 96-and 168 h. At each time point, 2 × 10 parasites /time point were transferred to Schneider’s Drosophila medium at 22°C to assess their revival capacity at different time points. **(H)** Representative ITS1 PCR image from DNA extracted from total parasite pellets harvested after exposure to macrophage growth conditions. PCR products were resolved on a 1% agarose gel electrophoresis. M: Ladder; L1-L3: LD PCR products from parasites inoculated in DMEM-10 and totally pelleted after 24-, 72-and 168 h; L4-L6: LD PCR products from parasites inoculated in RPMI-10 for 24-, 72-and 168 h; L7: 10^6^ LD DNA as positive control; L8-L10: LS PCR products from parasites inoculated in DMEM-10 and totally pelleted after 24-, 72-and 168 h; L11-L13: LS PCR products from parasites inoculated in RPMI-10 and totally pelleted after 24-, 72-and 168 h; L14: 10^6^ LS DNA as positive control (where L represents Lane number). **(I)** IL-12 (p40) cytokine level was measured by ELISA from cell supernatant infected with LD, LS and co-culture at 72 h.p.i. For (A-G and I), the data were represented as mean ± SD of a minimum of three independent experiments. Statistical comparisons were made using an unpaired two-tailed Student’s t-test or one way-Anova where p < 0.05 (*), p < 0.01 (**), p < 0.001 (***), p < 0.0001 (****), ns = not-significant.

To further assess species composition in co-infected samples, ITS1 band intensities were analysed using ImageJ. Despite higher initial LD abundance (10:1), the IC LD: LS ratio shifted to ∼1:150, 1:12 and 1:66 at 48-, 96 and 168 h.p.i time points respectively suggesting preferential amplification of LS over LD under co-infection scenario. Overall data supported active proliferation rather than persistence of LS inside macrophages alone or when present with LD over 168 h. Interestingly, ImageJ analysis demonstrated that LS replicated far more efficiently thereby rupturing macrophages and getting released in the extracellular milieu at higher proportions up to 333, ∼13 and 2500-fold at 48-, 96-and 168 h.p.i respectively, compared to LD in co-infected cells. Total parasite yield (EC + IC) except 48 h.p.i were always higher in co-infection conditions compared to mono-infection for all other time points tested. Perhaps, that is why, EC amastigote load was also higher for co-infection compared to mono-infections (except the 48 h time point) due to increased IC load of parasites in infected macrophages. (Fig. 1B), (Supplementary Table S1).

MTT analysis further proved that macrophage viability varied markedly across infection conditions. While LD caused a moderate reduction in viability (63.7% survival), LS infection led to a pronounced decrease (36% survival), indicating higher cytolysis. A similar low viability was observed in 2:1 co-infection (33.1% survival), consistent with LS-dominant effects right from the beginning. In contrast, increasing LD proportions during co-infection, progressively restored cell survival, with 5:1 and 10:1 co-infection showing high viability (83.4% and 90.4% survival, respectively), compared to LD or LS mono-infections thereby approaching cell intactness like uninfected levels (Fig. 1E).

#### 3.1.2. Temporal quantification revealed differential replication and population dynamics of LD and LS in co-infected human macrophages

Similar to findings in murine macrophages, both LD and LS successfully infected and replicated within differentiated human monocyte-derived macrophages (THP-1). Intracellular LS load significantly increased at 96-and 168 h.p.i. compared to 48 h.p.i by ∼2.7 and 10.8-fold respectively. In case of co-infection, the total intracellular parasite load also significantly increased over time at 96-and 168 h.p.i compared to 48 h.p.i by ∼2.1 and ∼ 4.9 -fold respectively. The progressive increase in intracellular LS burden supports robust parasite replication, rather than passive persistence within human macrophages irrespective of the presence of LD over the 168-h infection course. LD also exhibited a significant increase in intracellular burden at 168 h.p.i. compared to 48 h.p.i. by ∼2.1-fold (Fig. 1C) and (Supplementary Fig. S2).

ImageJ-based analysis of ITS1 PCR products indicated a temporal shift in species composition during co-infection. Inside macrophages (IC), the initial LD: LS ratio (10:1) increased at the beginning to 67:1 at 48 h.p.i, which later decreased to 13:1 and 1:1.3 at 96-and 168 h.p.i. respectively suggesting the preferential replication of LS over LD under co-infection scenario. Similar to murine macrophage cells, the total parasite yield (EC + IC) except 48 h.p.i were always higher in co-infection conditions compared to both mono-infections for all other time points tested. These results further validated the replication of monoxenous parasite LS inside mammalian macrophages. Interestingly, in co-infected cells, the LD: LS ratio in supernatant (EC) reversed from 2.5:1 at 48 h to 1:5 and 1:50 at 96-and 168 h.p.i. respectively. This suggested that the LS parasites were released rapidly from macrophages at later time points (upon infected cell rupture) which resulted in greater to at least equal LD: LS ratio inside macrophages from 48-to 168 h time points respectively (Supplementary Table S2). In THP-1 cells, unlike LD at 168 h.p.i., LS and co-infection conditions did not exhibit any significant change in the extracellular parasite burden over the tested time points (Fig. 1D).

Although the ratios are derived from semi-quantitative densitometric measurements, the consistent directional shift across replicates supports a progressive redistribution of LS during infection. These observations are indicative of differential population dynamics between LD and LS, rather than precise quantitative differences.

### 3.2. Macrophage internalized parasites transformed to promastigotes when cultured in Schneider’s Drosophila medium

#### 3.2.1. Parasites remained viable after infecting murine macrophage cells RAW 264.7

For every time point yield (72-168 h), transformation of amastigotes to promastigotes (reactivation) was carried out in sub-culture flasks containing supplemented Schneider’s Drosophila medium. These subcultures were examined at 48 h post-transfer under the microscope to check for any viable and motile parasites i.e. transformed promastigotes. Four representative microscopic fields were observed per flask and motile cells were enumerated (Supplementary Table S3 and Supplementary Fig. S3).

The 48 h counts upon transfer from each time-point harvest showed that the number of parasites increased progressively in the sub-culture flasks from earlier to later time-point harvests, indicating increasing amastigote production within macrophages over 168 h. Sub-cultures were further incubated, and two representative fields were observed per flask again at 11 days post-infection (d.p.i). Increased parasite counts at 11 d.p.i. in the subculture flasks compared to corresponding 48 h counts indicated successful replication of transformed promastigotes upon revival of amastigotes in supplemented Schneider’s Drosophila medium. For instance, LS subculture from the 120 h original harvest showed an increase in parasite count between early (Count: 23 ± 2; 2 days post subculture) and later observations (Count: 67 ± 17; 11 days subculture), confirming parasite viability and proliferation in Schneider’s Drosophila medium (Supplementary Table S4 and Supplementary Fig. S4). At 12 d.p.i., viable, motile cells in the total subcultures were also quantified using a haemocytometer (Fig. 1F and Supplementary Table S5), supporting the microscopic counts.

To compare the intrinsic growth properties of the revived parasites, 10^6^ viable parasites from the 12^th^ day harvest of the subculture flasks was further cultured and monitored up to 168 h in Schneider’s Drosophila medium. Haemocytometer counts showed that LS-containing cultures (alone or in mixtures) grew to higher titres at 96 h and 168 h compared to LD-only cultures (Supplementary Table S6 and Supplementary Fig. S5).

Importantly, additional observations revealed that adhered, non-phagocytosed parasites could remain static but viable in DMEM-10 up to 48 h during macrophage infection, and when transferred to Schneider’s Drosophila medium they regained the capacity to proliferate. However, such revival did not occur in case of later harvests of infected macrophages (>48 h). Non-internalised parasites could neither persist beyond 48 h in DMEM-10, nor regained viability when transferred to Schneider’s Drosophila medium at 22°C (Fig. 1G). This clarified that any apparent revival beyond 48 h observed in Schneider’s Drosophila medium, was only from intracellular amastigotes that transformed in the subculture flasks.

#### 3.2.2. LD or LS could not replicate to higher titres in DMEM-10 or RPMI-10 at 37°C

In order to test whether free parasites could replicate in macrophage culture conditions, viable LD and LS parasites were inoculated into 12-well plates at 10^6^ parasites /well and incubated at 37°C in DMEM-10 and RPMI-10 for 7 days. No motile or viable parasites were observed at the end of the experiment.

It was further evident from both molecular analysis (PCRs) as well as microscopic analysis neither LD nor LS could grow in DMEM-10 or RPMI-10 at 37°C when tested over a time point of 168 h. The titre was estimated (ImageJ analysis) to stay around 10^4^-10^5^ parasites per 5 mL (10^4^ for LD and 10^5^ for LS) culture over a period of 24-168 h post-inoculation in the aforesaid growth media (Fig. 1H).

As mentioned earlier, LS stayed static but viable up to 48 h but then became non-viable and could not be revived beyond 48 h of culture. Hence, it can be concluded that LS detected in the sub-cultures (in Schneider’s Drosophila medium at 22°C) from macrophage-infected yields at 72 h time point onwards, must had replicated inside the macrophages and revived as promastigotes in the subculture media.

### 3.3. IL-12 (p40) cytokine profiling reflected immune modulatory role of LD, LS and co-infective states

IL-12 (p40) ELISA from 72 h supernatant of infected RAW 264.7 cells revealed that LD-infected cells induced the highest IL-12 (p40) protein expression by approximately 2.62-fold compared to the uninfected control cells. LS-infected cells did not show any significant change in IL-12 (p40) levels compared to LD-infected cells. In contrast, co-culture-infected cells showed reduced IL-12 (p40) expression, approximately 1.43-fold lower compared to LD-infected cells (Fig. 1I). It may be an indication of dampened pro-inflammatory signalling, possibly linked to reduced macrophage activation, higher parasite load, and prolonged persistence.

### 3.4. Lepsey NLV1 was detected more inside mammalian macrophages compared to extracellular milieu

#### 3.4.1. Intracellular persistence of Lepsey NLV1 without temporal proliferation in murine macrophages

Quantitative RT-PCR analysis at 48 h.p.i. confirmed the presence of Lepsey NLV1 RNA within RAW 264.7 macrophages. Intracellular (IC) virus level was approximately 20-130-fold higher than the extracellular (EC) fraction (Fig 2A). Lepsey NLV1 titre (gE: RNA genome Equivalent) was further estimated from 48-168 h.p.i. in a time point assay. A significant increase in intracellular viral load was observed in LS-mono-infected cells, with a 1.2-fold elevation at 168 h.p.i. compared to 48 h.p.i. Although, significantly greater EC virus titre was detected in co-infection at 96 h.p.i compared to 48 h.p.i , no significant increase were detected in either intracellular viral titres or total viral titres (EC+IC) in the co-infected group at 96 h.p.i. and 168 h.p.i. relative to 48 h.p.i. IC virus titres were found to remain stable at (∼7.4-8.8 × 10^7^gE /well) in co-infection scenario, whereas EC titres were maintained at lower levels (5-6 × 10^6^ gE /well) (Figs. 2B, 2C) and (Supplementary Table S7). Collectively, these results suggested that while LS mono-infection supported progressive viral accumulation over time, co-infection was associated with a relatively stable viral replication profile throughout the infection period. LS carrying Lepsey NLV1 efficiently entered RAW 264.7 macrophages, confirming strong intracellular localization.

**Fig. 2.**
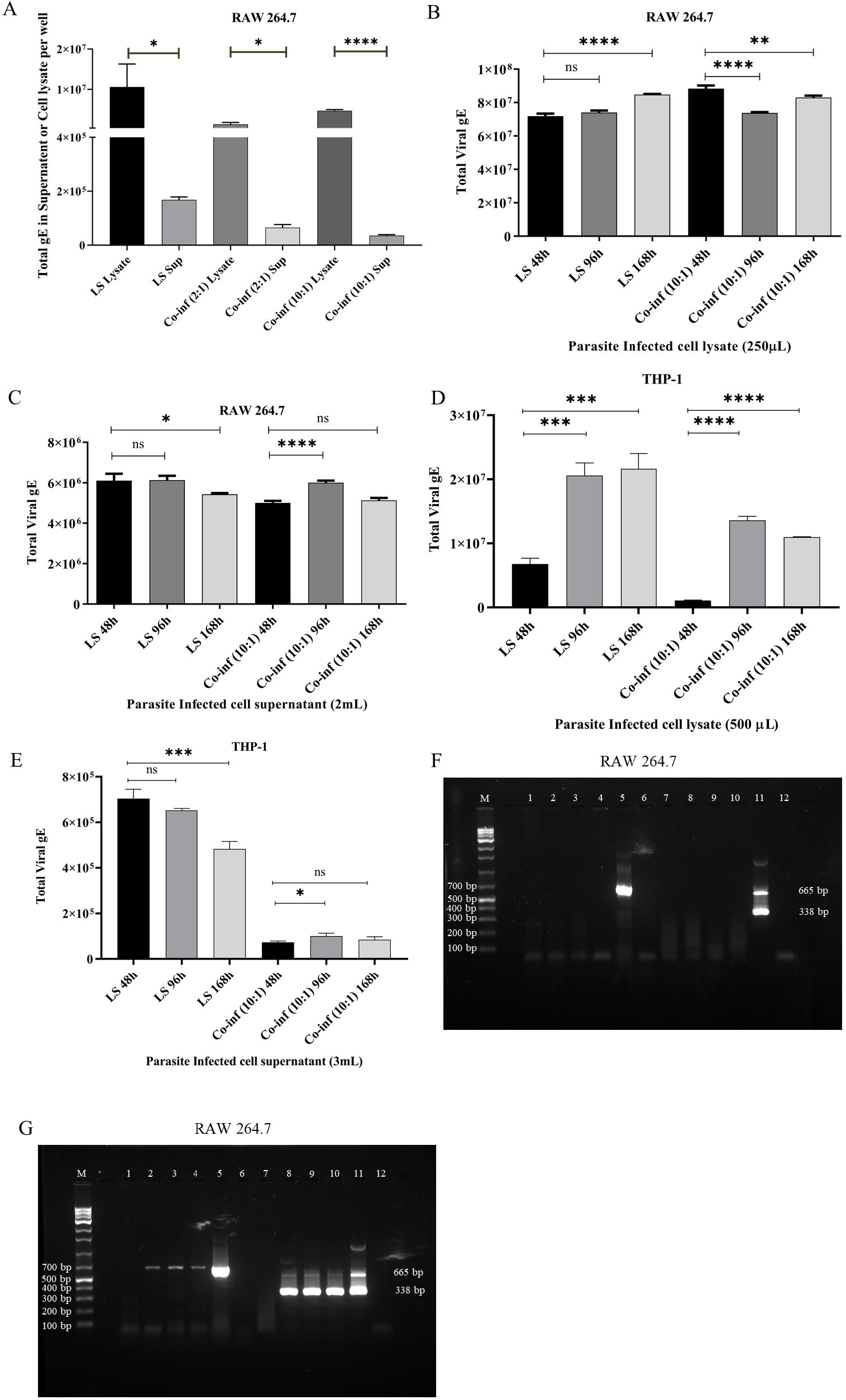
Quantification of virus load from parasite infected macrophage cells. Virus titres were assessed in culture **(A)** cell lysate and supernatant from parasite infected RAW 264.7 cells at 48 h.p.i. Virus load quantified from infected RAW 264.7 macrophage **(B)** cell lysate and **(C)** supernatant at 48-, 96-and 168-h.p.i. Quantification of virus titres from culture **(D)** cell lysate and **(E)** supernatant of parasite infected THP-1 cells at 48-, 96-and 168 h.p.i. Representative images of semi nested RT-PCR analysis for detection of *Leptomonas seymouri* narna-like virus 1 from virus infected RAW 264.7 **(F)** cell lysate and **(G)** supernatant at 48-, 72-and 96 h.p.i. M: marker; L1-L6: represents 1st round of PCR products and L7-L12: represents 2nd round of PCR products. L1 and L7: cell control; L2-L4 and L8-L10: 48-,72-and 96 h.p.i. samples respectively; L5 and L11: positive control, L6 and L12: negative control (where L represents lane number). For (A-E), the data are represented as mean ± SD of a minimum of three independent experiments. Statistical comparisons were made using an unpaired two-tailed Student’s t-test or one way-Anova where p < 0.05 (*), p < 0.01 (**), p < 0.001 (***), p < 0.0001 (****), ns = not-significant.

#### 3.4.2. Human monocyte-derived macrophages supported progressive intracellular replication of Lepsey NLV1 unlike murine macrophages

Similar analysis confirmed the presence of Lepsey NLV1 RNA in differentiated human monocyte-derived macrophages (THP-1) at multiple post-infection time points. Intracellular viral load in LS mono-infected cells increased significantly over time, exhibiting 3.1-and 3.3-fold higher titres at 96 h.p.i. and 168 h.p.i., respectively, compared to 48 h.p.i. Interestingly, in the co-infected group, intracellular viral load was significantly elevated, showing 12.7-and 10-fold increases at 96 h.p.i. and 168 h.p.i., respectively, relative to 48 h.p.i., suggesting better viral persistence in co-infection scenario compared to mono-infection (Fig. 2D).

No progressive increase was observed in extracellular virus titre for both LS-mono infection as well as co-infected cells at 168 h.p.i compared to 48 h.p.i. (Fig. 2E). The accumulation of virus predominantly within the intracellular compartment, coupled with relatively stable extracellular virus load, suggesting efficient virus replication within LS amastigotes with limited virus release in THP-1 derived human macrophage cells. Total virus load (EC + IC) significantly increased over time possibly due to increase in total LS count over time from 48-168 h.p.i (Supplementary Table S8). These finding indicate that THP-1 cells support sustained virus replication where Lepsey NLV1 persisted at **∼**10 –100 -fold higher levels intracellularly than in the extracellular environment.

#### 3.4.3. Lepsey NLV1 extracted from LS, could not alone infect macrophage cells

In contrast, purified Lepsey NLV1 (obtained by sonication, sedimentation and filtration of LS cultures) failed to infect RAW 264.7 macrophages. Virus RNA was detected from 48 to 96 h.p.i. It was limited to the supernatant and remained comparable to inoculum levels, with no intracellular detection (Figs. 2F and 2G). These findings indicated that purified Lepsey NLV1 alone could not enter or replicate in macrophages independently.

### 3.5 Microscopic observation of mammalian macrophages infected with LD or LS

Spherical shaped amastigote images were observed under microscope at 72 h.p.i. inside Giemsa stained macrophages. LS amastigotes were visible within both RAW 264.7 (Figs. 3A-C) and THP-1 cells (Figs. 3D-F), albeit less conspicuously compared to LD.

**Fig. 3:**
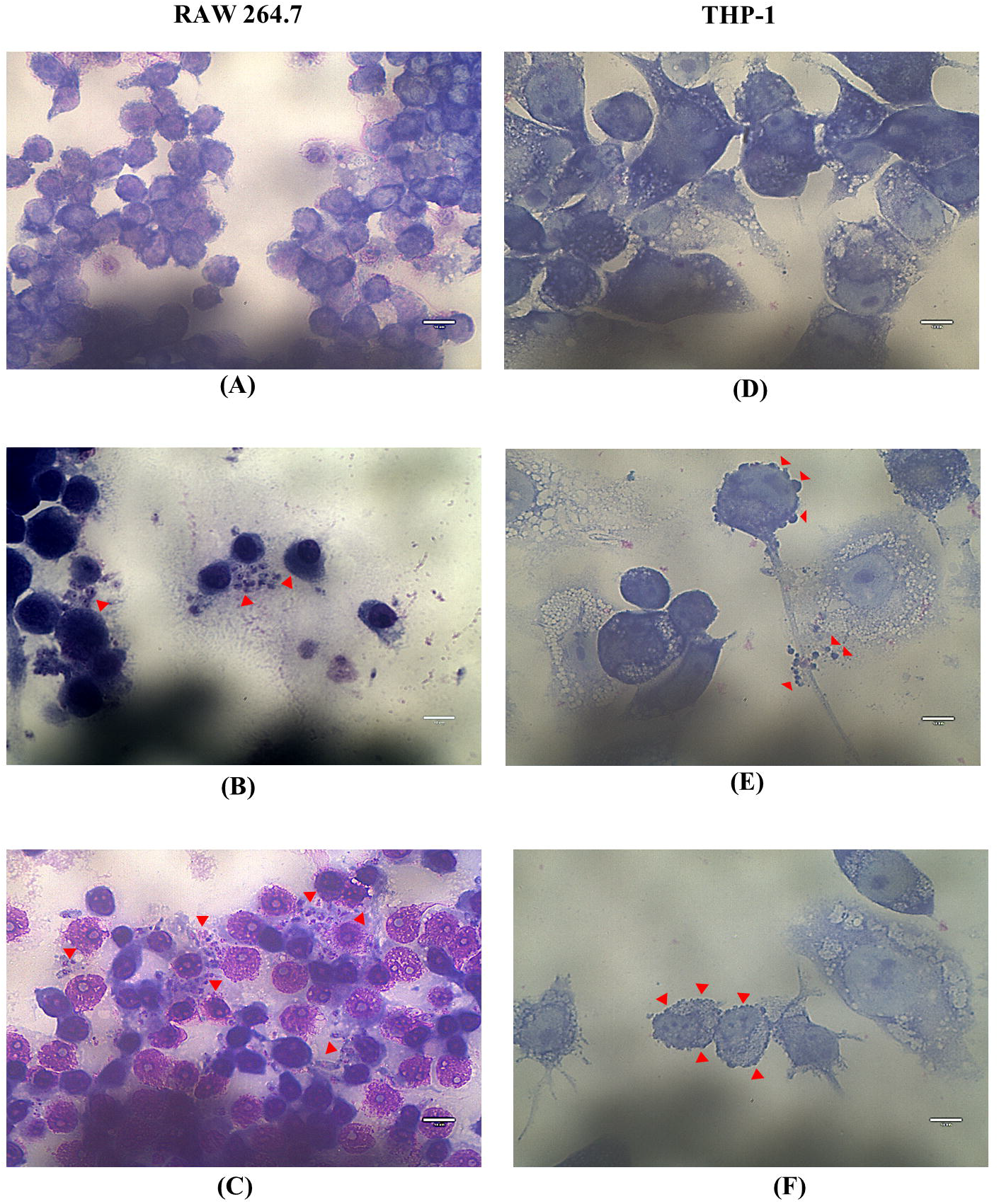
Microscopic examination of Giemsa-stained infected macrophage cells at 72 h.p.i. **(A)** uninfected RAW 264.7 macrophage cells; **(B)** RAW 264.7 macrophages infected with LD; **(C)** RAW 264.7 macrophages infected with LS; **(D)** Uninfected THP-1 macrophage cells; **(E)** THP-1 cells infected with LD; **(F)** THP-1 cells infected with LS. All images were acquired under a light microscope at 1000× magnification. Scale bar: 10 µm.

## Discussion

Here, we conducted a comprehensive assessment of the infectivity of LS, a parasite frequently co-isolated with LD from kala-azar affected individuals in India **(Sukla et al., 2017)**. *In vitro* results from this study, suggested that LS could infect and replicate in murine as well human macrophage cells such as RAW 264.7 and THP-1 respectively. Although the two parasites showed increased intracellular DNA over time in RAW 264.7, LS was more abundantly detected in intracellular (IC) and extracellular (EC) compartments, both alone and as co-infecting parasite with LD. In the co-infection setting, the progressively elevated intracellular burden of LS than LD points to highly efficient intracellular replication resulting in higher IC load. This resulted in the release of more LS amastigotes into the extracellular milieu even from few ruptured /dead infected macrophages (cell death was lowest in co-infection). This behaviour contrasts with the more controlled intracellular strategy of LD, which is known to preserve host cell integrity to sustain its niche **(Gupta et al., 2016)**. The coexistence of these opposing strategies suggests a dynamic interplay in which LS-driven cytolysis appears to promote parasite dissemination in the host body fluid (blood, lymph) thereby enhancing the possibility of parasite uptake by the vector. LD-mediated stabilization of macrophages limits excessive cell lysis and parasite dissemination. Notably, from MTT analysis, the survival of infected cells at higher LD: LS ratios during infection relative to LS mono-infection implies that LD can partially counterbalance LS-induced rupture, thereby maintaining a viable intracellular reservoir and long-term persistence of LS. Moreover, co-infection experiments revealed that the presence of LD did not adversely affect LS replication. On the contrary, in murine macrophage cells, LS replicated faster compared to LD resulting in 1:66 ratio in IC milieu at 168 h.p.i. despite 10-fold more abundance of LD in the original inoculum (LD: LS= 10:1). Relative to 48 h.p.i., intracellular LS burden increased more markedly during co-infection (4.4-and 7.3-fold at 96 and 168 h.p.i., respectively) than during LS mono-infection (3.1-and 5.3-fold), indicating enhanced intracellular persistence and proliferation of LS in the presence of LD.

Despite greater survival of macrophages, the EC parasite load was higher in case of co-infection suggesting greater IC parasite load per macrophage and thereby getting release in the extracellular milieu upon cell rupture. Thus, amastigote dissemination in case of co-infection was comparable to LS mono-infection despite lower macrophage rupture than LS mono-infected macrophages. A comparable earlier study, likewise, demonstrated critical preliminary evidence of the varying ability of LD strains to alter macrophage function, notably impairing the digestion and facilitating the survival of subsequently internalized *Leptomonas* sp. **(Kutish and Janovy, 1981).**

The THP-1 macrophage infection model further corroborated our observations in murine RAW 264.7 cells, confirming that LS could infect and replicate within differentiated human monocyte–derived macrophages. The intracellular scenario in THP-1 cells was similar to murine macrophage cells where IC parasite loads significantly increased over time for all the tested time points except at 96 h.p.i compared to 48 h.p.i for LD infected cells. Importantly, the increase in combined parasite load (EC+IC) over time further supported active replication of LS within a human macrophage environment, thereby extending our murine findings to a clinically relevant host cell system. The total parasite load was also highest for co-infection compared to both mono-infection from 96-to 168 h.p.i, thereby suggesting better parasite replication in the co-infected macrophages.

In co-infection, a marked temporal shift in LD: LS ratio was observed. While LD amastigote DNA predominated in the supernatant at 48 h.p.i. upon macrophage rupture, LS DNA became the dominant amastigote species in the extracellular medium at 96 and 168 h.p.i. This coincided with a sharp reduction in the intracellular LD: LS ratio. This reciprocal redistribution suggested progressive intracellular accumulation of LS followed by enhanced release into the extracellular milieu at later time points, likely due to higher parasite load in ruptured macrophages (similar to RAW 264.7). From overall findings, it was evident that LS replicated and persisted better over time under co-infection condition (∼10-195 fold) compared to mono-infection (∼2.6-10.4 fold) at the tested time points.

The rapid intracellular replication of LS, coupled with extensive host-cell rupture as evident from MTT assay, may provide a plausible explanation for why LS is generally not detected as a sole infectious agent in clinical settings. Excessive cytolysis could limit the establishment of a stable intracellular reservoir, thereby reducing the likelihood of long-term persistence and clinical detection. In contrast, during co-infection with LD, the parasite may benefit from the more controlled intracellular environment maintained by LD-infected macrophages, which preserves host-cell integrity and facilitates sustained parasite survival. Such a permissive niche could enable LS to persist for extended periods within the host, increasing the probability of its detection alongside LD in clinical samples. This interpretation is consistent with the frequent co-isolation of LS with LD from kala-azar and PKDL patients. Notably, LD and LS DNA remain detectable in PKDL skin biopsy samples even up to 7 years after apparent cure of visceral leishmaniasis, underscoring the long-term stability and clinical relevance of such co-infections **(Ghosh et al., 2012b)**.

Our observations didn’t align with a previous report that explained LS, as a monoxenous kinetoplastid, could not replicate in mammalian cells (although different macrophage cell lines, J774 and BMM were used) **(Kraeva et al., 2015).** Taken together, the RAW 264.7 and THP-1 infection models collectively supported a broader host adaptability of LS, highlighting its capacity for sustained intracellular replication, enhanced cyto-pathogenicity, and efficient dissemination from mammalian macrophages especially under co-infection conditions.

This is further consistent with earlier hypotheses that LS may act as a commensal or co-endosymbiont in LD infections, potentially influencing disease outcomes **(Srivastava et al., 2010)**. The viability assays demonstrated that parasites harvested from macrophage infections could be successfully revived in serum-supplemented Schneider’s Drosophila medium across multiple time points (72 h-168 h), confirming their persistence in intracellular environments. The progressive increase in parasite counts during sub-culturing underscored that LS retained their proliferative potential post-macrophage passage. However, the inability of free, non-internalised parasites to persist or replicate in macrophage culture conditions indicated that parasite survival beyond 48 h was exclusively attributable to an intracellular niche. Transfer of known number of parasites (2 x 10^6^) from macrophage culture media to Schneider’s Drosophila medium beyond 48 h up to 168 h.p.i. didn’t result in any restoration of motility of parasites even after 9 days incubation and the final yields came less than inoculum. Only static parasites at 48 h in macrophage media gained motility and replicated in Schneider’s Drosophila medium. These observations ruled out extracellular replication as a confounding factor and strengthened the conclusion that parasite revival from cultures beyond 72 h.p.i. represented genuine intracellular survival and release. Conversely, neither LS nor LD parasites were able to replicate in DMEM-10 or RPMI-10 at 37°C media used during macrophage infections where parasite counts remained static and no motile forms were detected up to 8 days post-inoculation. These data convincingly ruled out extracellular replication in culture medium, implicating intracellular macrophage environments as the primary niche for LS persistence and propagation during infection, consistent with previous suggestion of LS adaptation under host-induced selective pressures **(Srivastava et al., 2010; Ghosh et al., 2012a).**

Dampened IL-12 (p40) secretion upon co-infection compared to mono-infections suggested a possible evasion mechanism from host immune system, consistent with the frequent detection of LD-LS co-infection in clinical VL samples. These observations aligned with our previously published observation of reduced IL-18 levels in human serum samples in case of co-infection cases compared to LD only samples **(Sukla et al., 2022)**. Such permissive environment also supports the enhanced replication and persistence of parasites under co-infection setting. Taken together, these results emphasized the importance of the host cellular environment in sustaining parasite viability. It suggested that LS possessed enhanced capacity to endure intracellular stress especially in co-infection scenario, a feature that might redefine its ecological persistence and pathogenic potential.

Microscopic evaluation of Giemsa-stained RAW 264.7 and THP-1 cells revealed amastigote-like forms of both LD and LS at 72 h.p.i. LD-infected cells exhibited more distinct amastigote forms. In contrast the presence of LS in intracellular compartments, albeit less prominently, suggested a morphologically adaptive, possibly amastigote-like transformation of LS under intracellular stress. The persistence of LS in mammalian macrophage cells further supported our molecular and growth-based observations of transient intracellular survival.

This study substantiated that LS, carrying Lepsey NLV1, could enter and persist within mammalian macrophages. Quantitative RT-PCR revealed higher intracellular virus loads than extracellular milieu at 48 h.p.i. This indicated efficient LS uptake and potential virus retention or amplification inside macrophages. Detected intracellular titres (∼10^6^-10^7^ gEs) contrasted with (∼10^5^-10^6^ gEs) in supernatant. This indicated robust intracellular localisation. This pattern remained relatively stable over 168 h in murine macrophages, implying a tightly controlled virus state inside macrophage cells. The comparable titres post-infection possibly reflected macrophage-mediated restriction or a steady-state balance of replication and degradation. Interestingly, in THP-1, intracellular virus loads increased ∼12.7-and 10-fold in co-infection compared to 3.1-and 3.3-fold increase in LS mono-infection at 96 and 168 h.p.i. relative to 48 h.p.i. These higher virus titres are probably due to more active replication of LS in permissive, surviving macrophages under co-infection scenario compared to LS mono-infection. The overall parasite and virus infection dynamics are represented with a schematic highlight (Fig. 4). Notably, purified Lepsey NLV1 could not enter or replicate in macrophages independently, thereby highlighting LS as a necessary carrier, likely delivering the virus via phagocytosis of infected parasites rather than direct cellular entry. Recent LD cases in Himachal Pradesh reported LS co-infection, raising questions about the virus’s role in atypical cutaneous manifestations **(Thakur et al., 2020).** The exact impact of Lepsey NLV1 dynamics and host-pathogen interaction, however, remained to be fully explored.

**Fig. 4:**
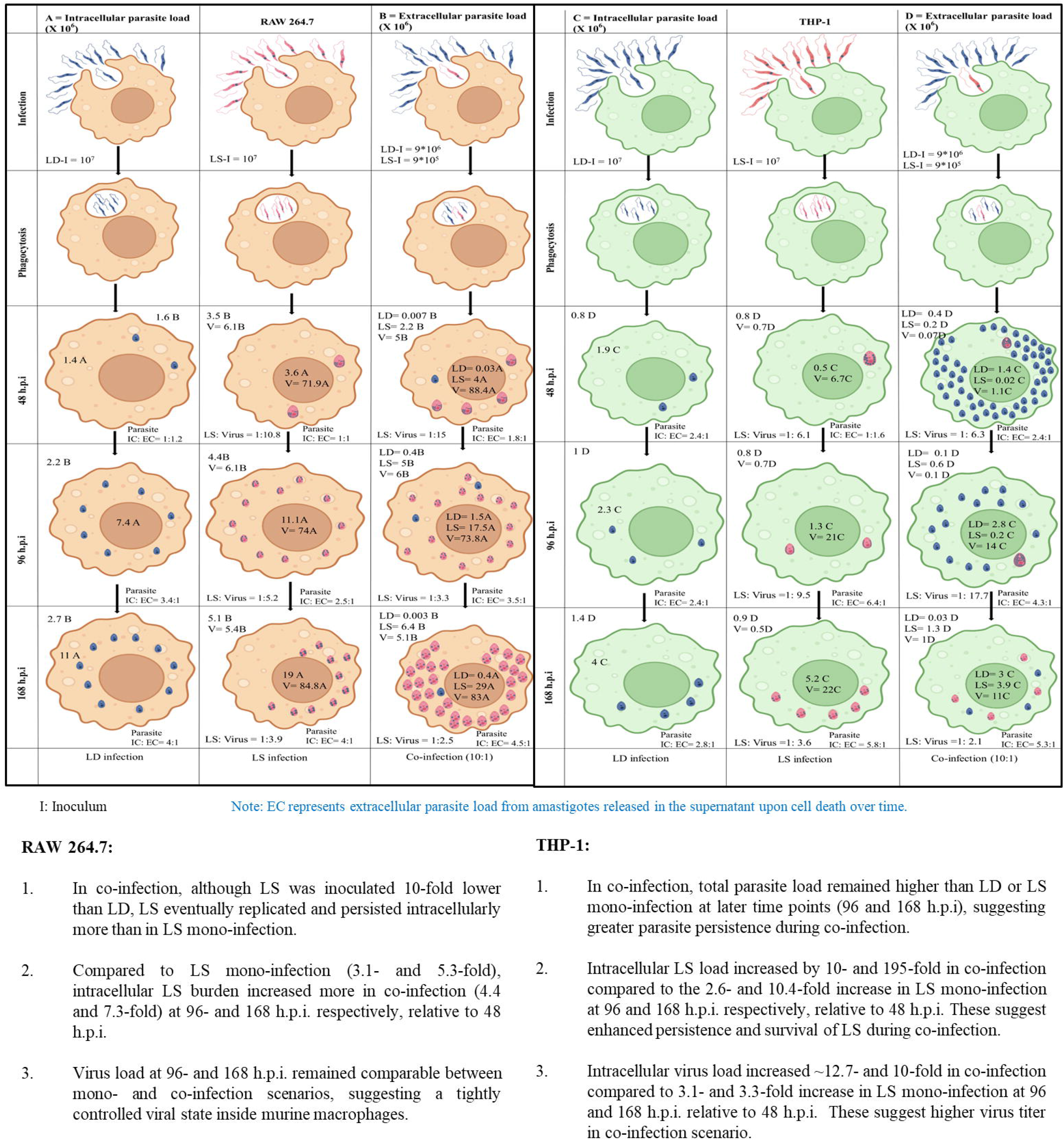
Schematic representation of parasite and virus load dynamics during mono-infection and co-infection in murine and human macrophage models. **(A)** Intracellular (IC) parasite load in RAW 264.7 macrophages, expressed as parasite DNA count (×10^6^), at indicated time points (48-, 96-and 168-hours post-infection (h.p.i.)). **(B)** Extracellular (EC) parasite load in the corresponding culture supernatant of RAW 264.7 macrophages, expressed as parasite DNA count (×10^6^), at 48-, 96-and 168 h.p.i. **(C)** Intracellular (IC) parasite load in THP-1-derived human macrophages, expressed as parasite DNA count (×10^6^), at 48-, 96-and 168 h.p.i. **(D)** Extracellular (EC) parasite load in the corresponding culture supernatant of THP-1-derived macrophages, expressed as parasite DNA count (×10^6^), at 48-, 96-and 168 h.p.i. Data represent mono-infection with *Leishmania donovani* (LD), mono-infection with *Leptomonas seymouri* (LS), and co-infection (LD: LS at 10:1 ratio).

Taken together, our findings presented a thought-provoking re-evaluation of LS, particularly in co-infection scenarios with LD. This study critically redefined the biological and pathogenic profile of LS, challenging its long-held classification as a non-infective, monoxenous parasite, but as a potentially replicative parasite capable of persisting within mammalian macrophages. The association of LS with the Lepsey NLV1 virus introduced a significant paradigm shift, highlighting a complex LD-LS-virus (triple pathogen) biology. It is noteworthy that *Leptomonas* sp. had been previously associated with infections in immunosuppressed patients **(Pacheco et al., 1998)**. These findings underscored the underestimated pathogenic potential of LS. It warrants a detailed mechanistic study into host-pathogen dynamics and its influence on *Leishmania* disease outcomes, particularly in co-infection contexts.

## FUNDING

The research was funded by Indian Council of Medical Research; grant number: Discovery/IIRP/SG-0959/2023 (GAP-473). The grant was given to S.B. The funders had no role in the study design, analysis and interpretation of data; in the writing of the manuscript; and in the decision to submit the manuscript for publication.

## AUTHORS’ CONTRIBUTION

S.B., S.D. and P.D.S. conceived and designed the experiments. S.D. and P.D.S performed the experiments equally. S.D. and P.D.S. performed data analysis and S.B. performed critical analysis of the data. S.B. provided funding, reagents, materials and analysis tools. S.B., S.D. and P.D.S. jointly wrote the original draft of the manuscript. All authors critically reviewed and modified and agreed on the current version of the manuscript.

## Supporting information

...but to lower counts compared to LS mono- or co-infection. (Fig. 1A) and (Supplementary Fig. S1).

## ACKNOWLEDGEMENT

Authors acknowledge the support received from the Director of CSIR IICB. S.B. also acknowledges AcSIR for support. S.D. and P.D.S acknowledges the support of UGC for their UGC-Research Fellowships. The authors would also like to acknowledge the use of relevant facilities at CSIR-IICB including the Central Instrumentation Facility (CIF). The authors acknowledge CSIR-IICB for providing all laboratory facilities for conducting the current work.

## DECLARATION OF INTEREST

S.B. reports financial support, article publishing charges, and travel were provided by Indian Council of Medical Research. The other authors declare that they have no known competing financial interests or personal relationships that could have appeared to influence the work reported in this paper.

## DATA AVAILABILITY

Further information and resource requests should be directed to and will be fulfilled by the corresponding author, Dr. Subhajit Biswas.

## Supporting Information Legends

**Fig. S1. Quantification of parasite load from LD, LS and co-infected (10:1) murine macrophage RAW 264.7 cells.** The graph represents the parasite load in both cell lysate (250 µL) and supernatant (2 mL) from LD, LS and co-infected (10:1) RAW 264.7 cells at 48-, 96-and 168-hour post-infection (h.p.i). Data are presented as mean ± SD from a minimum of three independent experiments. Statistical comparisons were made using one way-Anova, where p < 0.05 (*), p < 0.01 (**), p < 0.0001 (****), ns = not-significant.

**Table S1. Quantitative analysis of ITS1 qPCR-derived parasite load in RAW 264.7 cells**

Parasite load was quantified from supernatant and lysate samples collected at 48-, 96-and 168 h.p.i. The LD: LS ratio was determined using ImageJ by measuring the relative band intensities of ITS1 PCR products obtained from both supernatant and lysate at 48-, 96-and 168-h.p.i. Data are presented as mean ± SD from a minimum of three independent experiments.

**Table S2. Quantitative analysis of parasite load from infected THP-1 cells.**

Parasite load was quantified from supernatant and lysate samples collected at 48-, 96-and 168 h.p.i. The LD: LS ratio was determined using ImageJ by measuring the relative band intensities of ITS1 PCR products obtained from both supernatant and lysate at 48-, 96-and 168 h.p.i. Data are presented as mean ± SD from a minimum of three independent experiments.

**Fig. S2. Quantification of parasite load from LD, LS and co-infected (10:1) human macrophage THP-1 cells.**

The graph represents the parasite load in both cell lysate (500 µL) and supernatant (3 mL) from LD, LS and co-infected (10:1) THP-1 cells at 48-, 96-and 168-h.p.i. Data are presented as mean ± SD from a minimum of three independent experiments. Statistical comparisons were made using one way-Anova, where p < 0.05 (*), p < 0.01 (**), p < 0.001 (***), p < 0.0001 (****).

**Table S3. Representative counts of motile parasites per four microscopic fields at 48 h in subculture flasks.**

Motile parasites were counted in four representative microscopic fields following transfer of harvested cells to subculture flasks at respective time points. Data are presented as mean ± SD from a minimum of three independent experiments.

**Fig. S3. Graphical representation of motile parasite counts in subculture flasks at 48 h.**

The graph represents motile parasites counted in four representative microscopic fields following transfer of harvested cells to subculture flasks at respective time points. Microscopic examination was performed at 400X magnification. Data are presented as mean ± SD from a minimum of three independent experiments.

**Table S4. Representative microscopic counts of motile parasites in subculture flasks at different time points.**

Representative parasite counts from two different microscopic fields were recorded in the same “different time points” subculture flasks at 11 days post-infection (d.p.i). Microscopic examination was performed at 400X magnification. Data are presented as mean ± SD from a minimum of three independent experiments.

**Fig S4. Graphical representation of motile parasite counts in subculture flasks at different time points.**

Motile parasites in the same “different time points” subculture flasks were counted using light microscopy at 11 d.p.i. Microscopic examination was performed at 400X magnification. Data are presented as mean ± SD from a minimum of three independent experiments.

**Table S5. Representative counts of motile and transformed parasites in subculture flasks at different time points -12 d.p.i.**

Motile and transformed parasites were quantified using a haemocytometer following transfer of harvested cells into subculture flasks at the indicated time points. Data are presented as mean ± SD from a minimum of three independent experiments.

**Table S6. Viable and motile parasite counts determined using a hemocytometer after 7 days of subculture**

Parasite cultures were initiated with an inoculum of 1 × 10 parasites for each condition using parasites harvested 12 d.p.i. Viable and motile parasites were counted using a hemocytometer based on active movement observed under light microscopy. Data are presented as mean ± SD from at least three independent experiments.

**Fig. S5. Graphical representation of viable and motile parasite counts after 7 days of subculture**

Parasite cultures were initiated with an inoculum of 1 × 10 parasites for each condition using parasites harvested 12 d.p.i. Viable and motile parasites were quantified using a hemocytometerbased on active movement under light microscopy at 100X magnification. Data are presented as mean ± SD from at least three independent experiments.

**Table S7. Quantitative analysis of viral load in infected RAW 264.7 macrophages**

Viral load was quantified from infected RAW 264.7 cell culture supernatant and cell lysate at 48-, 96-and 168 h.p.i. Data are presented as mean ± SD from a minimum of three independent experiments.

**Table S8. Quantitative analysis of viral load in infected THP-1 macrophages**

Viral load was quantified from infected THP-1 cell culture supernatant and cell lysate at 48-, 96-and 168 h.p.i. Data are presented as mean ± SD from a minimum of three independent experiments.

## Notes

### Summary of Updates

This version is based on improved analysis of the data.

