## Supplementary material for "Co-infection with *Leptomonas seymouri* enhances macrophage survival and promotes intracellular parasite persistence during *Leishmania donovani* infection": ...but to lower counts compared to LS mono- or co-infection. (Fig. 1A) and (Supplementary Fig. S1).

### Supplementary Data

**Fig. S1. Quantification of parasite load from LD, LS and co-infected (10:1) murine macrophage RAW 264.7 cells.**

The graph represents the parasite load in both cell lysate (250 µL) and supernatant (2 mL) from LD, LS and co-infected (10:1) RAW 264.7 cells at 48-, 96- and 168-hour post-infection (h.p.i). Data are presented as mean ± SD from a minimum of three independent experiments. Statistical comparisons were made using unpaired student's t-test, where $p < 0.05$ (*), $p < 0.01$ (**), $p < 0.0001$ (****), ns = not- significant.

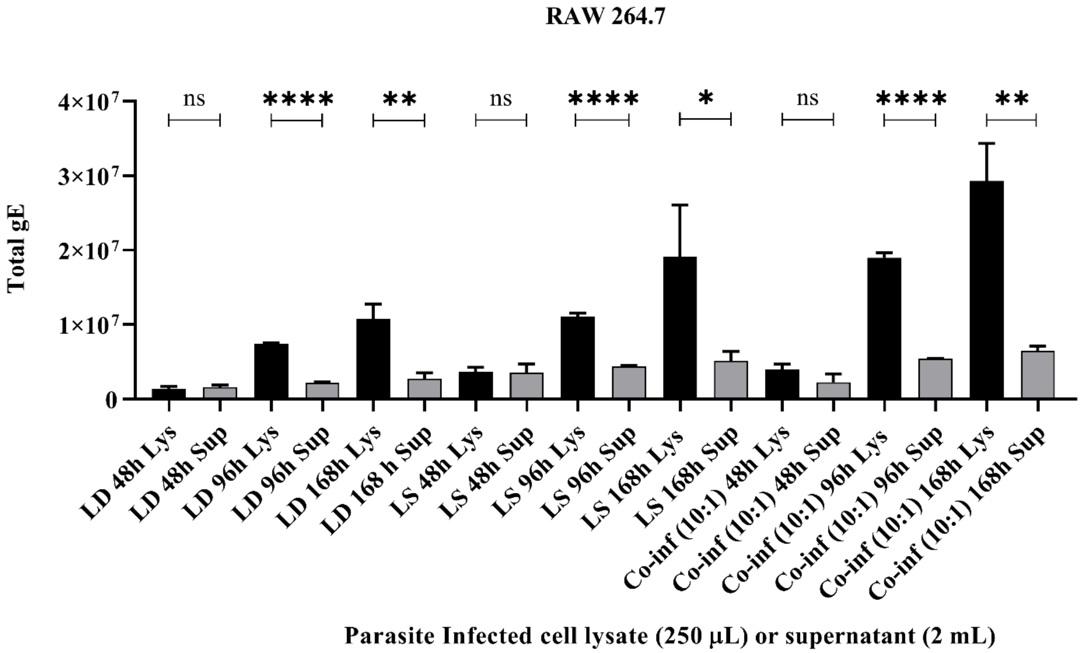

**Table S1. Quantitative analysis of ITS1 qPCR-derived parasite load in RAW 264.7 cells**

Parasite load was quantified from supernatant and lysate samples collected at 48, 96- and 168 h.p.i. The LD: LS ratio was determined using ImageJ by measuring the relative band intensities of ITS1 PCR products obtained from both supernatant and lysate at 48, 96- and 168-h.p.i. Data are presented as mean ± SD from a minimum of three independent experiments.

| SAMPLE | DNA copies/2 mL sup | Ratio (LD: LS) | DNA copies/250 µL lysate | Ratio (LD: LS) | Total Parasite DNA (EC+IC) | IC: EC |
|---|---|---|---|---|---|---|
| LD 48 h | 1.6 (± 0.27) x $10^6$ | | 1.4 (± 0.3) x $10^6$ | | 3 (± 0.6) x $10^6$ | 1:1.2 |
| LS 48 h | 3.5 (± 1.1) x $10^6$ | | 3.6 (± 0.6) x $10^6$ | | 7.2 (± 1.6) x $10^6$ | 1:1 |
| Co-inf (10:1) 48 h | LD- 6.6 (± 3.3) x $10^3$<br>LS- 2.2 (± 1.1) x $10^6$ | 1:333 | LD- 2.6 (± 0.5) x $10^4$<br>LS- 4 (± 0.7) x $10^6$ | 1:150 | 6.2 (± 1.8) x $10^6$ | 1.8: 1 |

| | | | | | | |
|---|---|---|---|---|---|---|
| LD 96 h | 2.2 (± 0.1) × $10^6$ | | 7.4 (± 0.1) × $10^6$ | | 9.6 (± 0.14) × $10^6$ | 3.4:1 |
| LS 96 h | 4.4 (± 0.1) × $10^6$ | | 1.1 (± 0.4) × $10^7$ | | 1.6 (± 0.41) × $10^7$ | 2.5:1 |
| Co-inf (10:1) 96 h | LD- 0.4 × $10^6$<br>LS- 5 × $10^6$ | 1:12.9 | LD- 1.5 (± 0.1) × $10^6$<br>LS- 1.8 (± 0.6) × $10^7$ | 1:12 | 2.4 (± 0.7) × $10^7$ | 3.5:1 |
| LD 168 h | 2.7 (± 0.8) x $10^6$ | | 1.1 (± 0.2) x $10^7$ | | 1.3 (± 0.3) x $10^7$ | 4: 1 |
| LS 168 h | 5.1 (± 1.2) x $10^6$ | | 1.9 (± 0.7) x $10^7$ | | 2.3 (± 0.6) x $10^7$ | 4: 1 |
| Co-inf (10:1) 168 h | LD- 2.6 (± 0.3) x $10^3$<br>LS- 6.4 (± 0.7) x $10^6$ | 1:2500 | LD- 4.3 (± 0.7) x $10^5$<br>LS- 2.9 (± 0.5) x $10^7$ | 1:66 | 3.5 (± 0.6) x $10^7$ | 4.5: 1 |

**Fig. S2. Quantification of parasite load from LD, LS and co-infected (10:1) human macrophage THP-1 cells.**

The graph represents the parasite load in both cell lysate (500 µL) and supernatant (3 mL) from LD, LS and co-infected (10:1) THP-1 cells at 48-, 96- and 168- h.p.i. Data are presented as mean ± SD from a minimum of three independent experiments. Statistical comparisons were made using unpaired student's t-test, where $p < 0.05$ (*), $p < 0.01$ (**), $p < 0.001$ (***), $p < 0.0001$ (****).

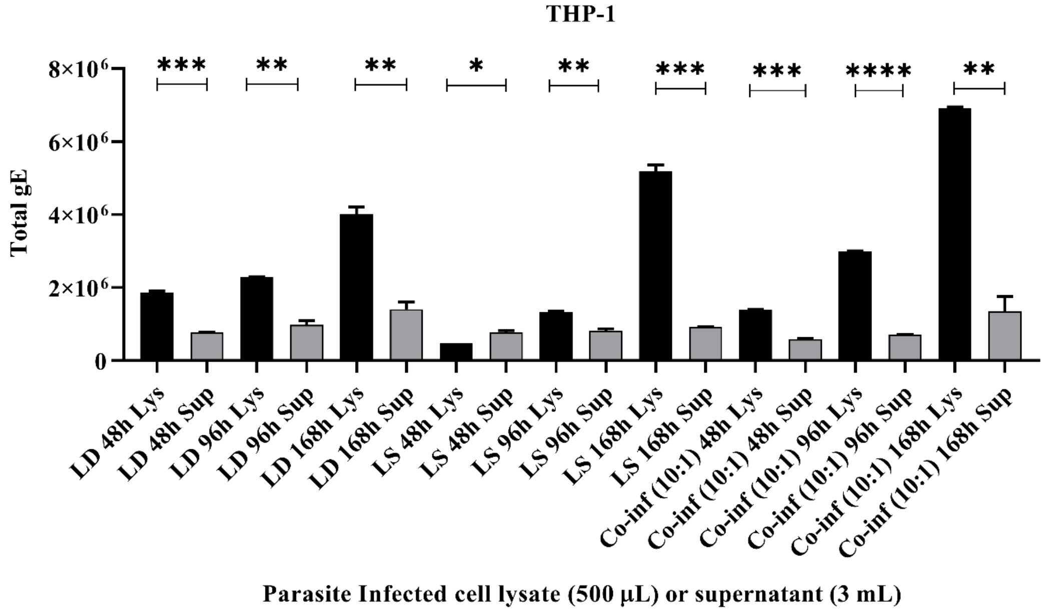

**Table S2. Quantitative analysis of parasite load from infected THP-1 cells.**

Parasite load was quantified from supernatant and lysate samples collected at 48, 96, and 168 h.p.i. The LD: LS ratio was determined using ImageJ by measuring the relative band intensities of ITS1 PCR products obtained from both supernatant and lysate at 48, 96, and 168 h.p.i. Data are presented as mean ± SD from a minimum of three independent experiments.

| SAMPLE | Parasite Load /3 mL sup | Ratio (LD: LS) | Parasite Load /500 µL lysate | Ratio (LD: LS) | Total Parasite Load (EC+IC) | IC: EC |
|---|---|---|---|---|---|---|
| LD 48 h | 7.6 (± 0.1) x$10^5$ | | 1.9 (± 0.04) x $10^6$ | | 2.6 (± 0.05) x $10^6$ | 2.4: 1 |
| LS 48 h | 7.6 (± 0.6) x $10^5$ | | 4.8 (± 0.003) x $10^5$ | | 1.2 (± 0.06) x $10^6$ | 1: 1.6 |
| Co-inf (10:1) 48 h | LD- 4.2 (± 0.1) x $10^5$ / LS- 1.7 (± 0.03) x $10^5$ | 2.5:1 | LD- 1.4 (± 0.01) x $10^6$ / LS- 2 (± 0.02) x $10^4$ | 67:1 | 1.9 (± 0.02) x $10^6$ | 2.4:1 |

| | | | | | | |
|---|---|---|---|---|---|---|
| LD 96 h | 9.7 (± 1.1) x $10^5$ | | 2.3 (± 0.02) x $10^6$ | | 3.3 (± 0.1) x $10^6$ | 2.4: 1 |
| LS 96 h | 8.1 (± 0.5) x $10^5$ | | 1.3 (± 0.02) x $10^6$ | | 2.2 (± 0.08) x $10^6$ | 6.4: 1 |
| Co-inf (10:1) 96 h | LD- 1.2 (± 0.02) x$10^5$<br>LS- 5.8 (± 0.12) x$10^5$ | 1:5 | LD- 2.8 (± 0.004) x $10^6$<br>LS- 2.1 (± 0.003) x $10^5$ | 13:1 | 3.7 (± 0.02) x $10^6$ | 4.3: 1 |
| LD 168 h | 1.4 (± 0.2) x $10^6$ | | 4.0 (± 0.2) x $10^6$ | | 5.40 (± 0.002) x $10^6$ | 2.8: 1 |
| LS 168 h | 9.1 (± 0.1) x$10^5$ | | 5.19 (± 0.16) x $10^6$ | | 6.10 (± 0.17) x$10^6$ | 5.8: 1 |
| Co-inf (10:1) 168 h | LD- 2.6 (± 0.8) x $10^4$<br>LS- 1.3 (± 0.4) x $10^6$ | 1:50 | LD- 3 (± 0.02) x $10^6$<br>LS- 3.9 (± 0.02) x $10^6$ | 1:1.3 | 8.20 (± 0.43) x$10^6$ | 5.3: 1 |

**Table S3. Representative counts of motile parasites per four microscopic fields at 48 h in subculture flasks.**

Motile parasites were counted in four representative microscopic fields at 48 h following transfer of harvested cells to subculture flasks at respective time points (72 - 168 h). Data are presented as mean ± SD from a minimum of three independent experiments.

| PARASITE | No. of cells in 4 different fields in (72 hours) flask | No. of cells in 4 different fields in (96 hours) flask | No. of cells in 4 different fields in (120 hours) flask | No. of cells in 4 different fields in (144 hours) flask | No. of cells in 4 different fields in (168 hours) flask |
|---|---|---|---|---|---|
| LD | 6 ± 1 | 19 ± 1 | 35 ± 3 | 33 ± 2 | 36 ± 4 |
| LS | 5 ± 1 | 15 ± 1 | 23 ± 2 | 29 ± 1 | 30 ± 3 |
| Co-inf (2:1) | 11 ± 1 | 17 ± 1 | 20 ± 4 | 29 ± 2 | 30 ± 2 |
| Co-inf (10:1) | 13 ± 1 | 16 ± 1 | 25 ± 3 | 30 ± 4 | 34 ± 3 |

**Fig. S3. Graphical representation of motile parasite counts in subculture flasks at 48 h.**

The graph represents motile parasites counted in four representative microscopic fields following transfer of harvested cells to subculture flasks at respective time points (72 - 168 h). Microscopic examination was performed at 400X magnification. Data are presented as mean ± SD from a minimum of three independent experiments.

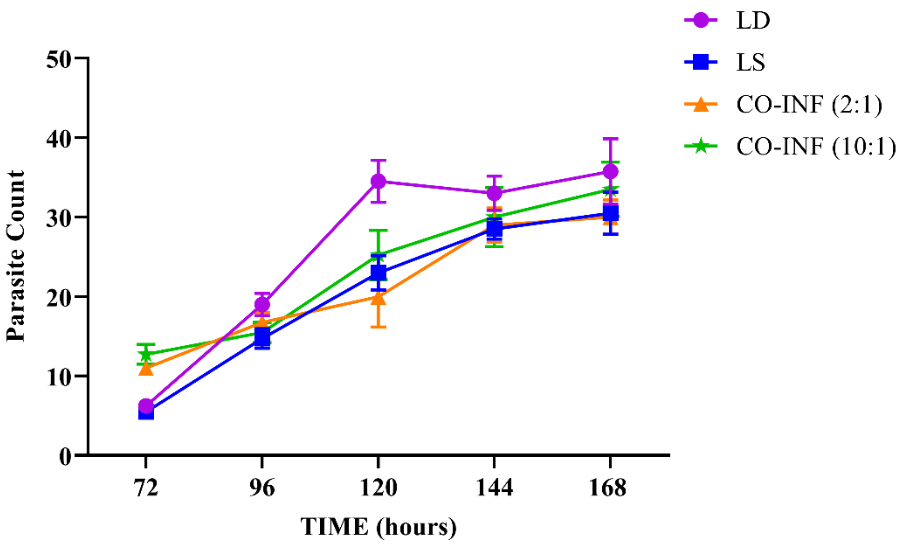

**Table S4. Representative microscopic counts of motile parasites in subculture flasks at different time points.**

Representative parasite counts from two different microscopic fields were recorded in the same "different time points" subculture flasks at 11 days post-infection (d.p.i.). Microscopic examination was performed at 400X magnification. Data are presented as mean ± SD from a minimum of three independent experiments.

| PARASITE | 72 hours flasks | 96 hours flasks | 120 hours flasks | 144 hours flasks | 168 hours flasks |
|---|---|---|---|---|---|
| LD | 52 ± 5 | 59 ± 4 | 71 ± 4 | 68 ± 8 | 78 ± 6 |
| LS | 55 ± 11 | 48 ± 7 | 67 ± 17 | 75 ± 8 | 70 ± 8 |
| Co-inf (2:1) | 62 ± 13 | 42 ± 6 | 77 ± 13 | 71 ± 3 | 62 ± 13 |
| Co-inf (10:1) | 55 ± 6 | 47 ± 13 | 80 ± 11 | 81 ± 5 | 79 ± 4 |

**Fig S4. Graphical representation of motile parasite counts in subculture flasks at different time points.**

Motile parasites in the same “different time points” subculture flasks were counted using light microscopy at 11 d.p.i. Microscopic examination was performed at 400X magnification. Data are presented as mean ± SD from a minimum of three independent experiments.

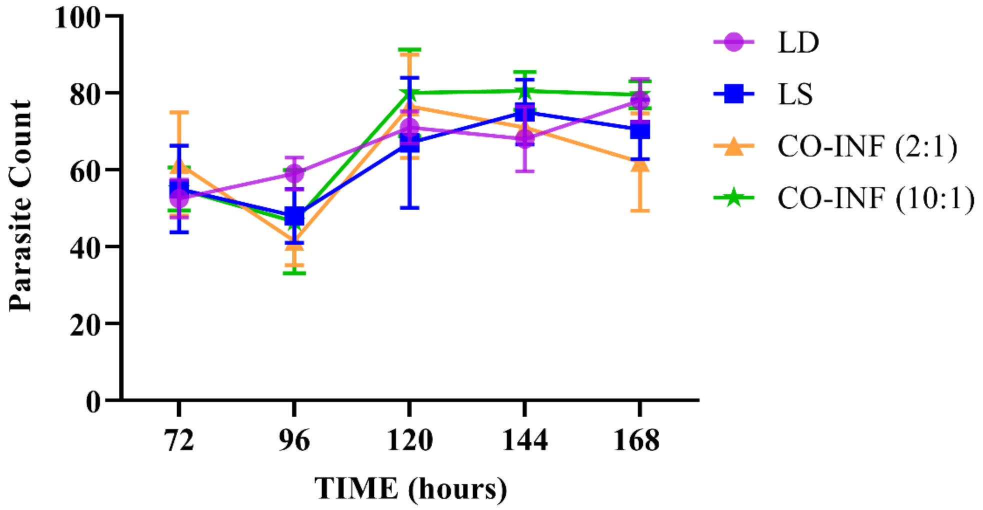

**Table S5. Representative counts of motile and transformed parasites in subculture flasks at different time points -12** d.p.i**.** Motile and transformed parasites were quantified using a hemocytometer following transfer of harvested cells into subculture flasks at the indicated time points. Data are presented as mean ± SD from a minimum of three independent experiments.

| PARASITE | 96 hours flask ($x10^7$)/ml | 168 hours flask ($x10^7$)/ml (x N = fold change w.r.t 96 h count) |
|---|---|---|
| LD | 5.4 ± 0.35 | 10.4 ± 0.56 (x 1.9) |
| LS | 4.9 ± 0.42 | 9.9 ± 0.21 (x 2) |
| Co-inf (2:1) | 5.5 ± 0.35 | 8.4 ± 0.28 (x 1.5) |
| Co-inf (10:1) | 6.4 ± 0.42 | 12.7 ± 0.14 (x 2) |

**Table S6. Viable and motile parasite counts determined using a hemocytometer after 7 days of subculture**

Parasite cultures were initiated with an inoculum of $1 \times 10^6$ parasites for each condition using parasites harvested 12 d.p.i. Viable and motile parasites were counted using a hemocytometer based on active movement observed under light microscopy. Data are presented as mean ± SD from at least three independent experiments.

| PARASITE | 96 hours flasks ($x10^7$)/ml | 168 hours flasks ($x10^7$)/ml (x N = fold change w.r.t initial inoculum) |
|---|---|---|
| LD | 2.3 ± 0.56 | 6.7 ± 0.42 (x 67) |
| LS | 5.4 ± 0.42 | 8.6 ± 0.28 (x 86) |
| Co-inf (2:1) | 6.9 ± 0.70 | 9.4 ± 0.14 (x 94) |
| Co-inf (10:1) | 6.1 ± 0.14 | 9.3 ± 0.14 (x 93) |

**Fig. S5. Graphical representation of viable and motile parasite counts after 7 days of subculture**

Parasite cultures were initiated with an inoculum of 1 × $10^6$ parasites for each condition using parasites harvested 12 d.p.i. Viable and motile parasites were quantified using a hemocytometer based on active movement under light microscopy at 100X magnification. Data are presented as mean ± SD from at least three independent experiments.

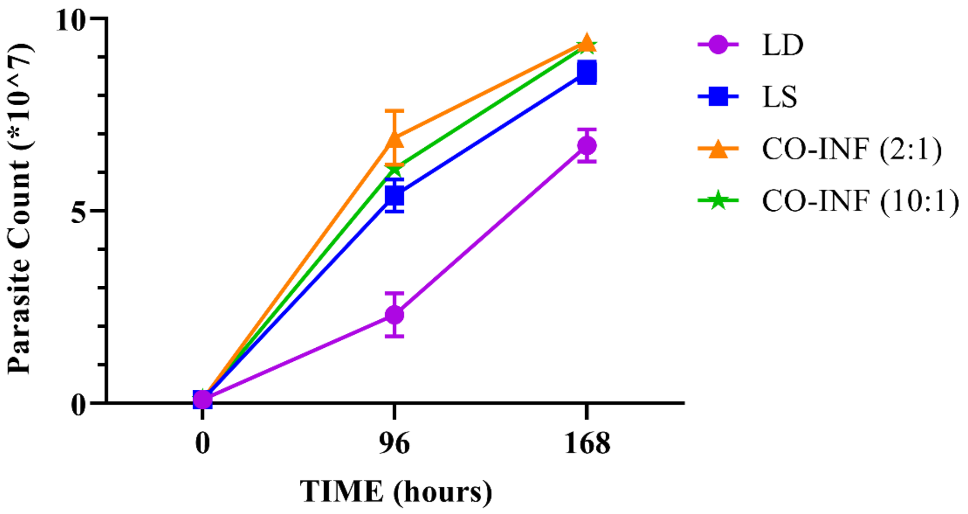

**Table S7. Quantitative analysis of viral load in infected RAW 264.7 macrophages**

Viral load was quantified from infected RAW 264.7 cell culture supernatant and cell lysate at 48, 96 and 168 h.p.i.. Data are presented as mean ± SD from a minimum of three independent experiments.

| Sample | Virus copies/2 mL sup | Virus copies/250 µL lysate | Total Virus gE (EC+IC) | Parasite: Virus Ratio | Virus Fold Increase Compared to only LS | IC: EC |
|---|---|---|---|---|---|---|
| LS 48 h | $(6.1 \pm 0.3) \times 10^6$ | $(7.2 \pm 1.5) \times 10^7$ | $(7.8 \pm 1.7) \times 10^7$ | 1:10.8 | | 11.8:1 |
| Co-inf (10:1) 48 h | $(5.0 \pm 0.1) \times 10^6$ | $(8.8 \pm 1.8) \times 10^7$ | $(9.3 \pm 1.8) \times 10^7$ | 1:15 | 1.4 | 17.7:1 |
| LS 96 h | $(6.1 \pm 0.2) \times 10^6$ | $(7.4 \pm 1.1) \times 10^7$ | $(8 \pm 1.3) \times 10^7$ | 1:5.2 | | 12.1:1 |
| Co-inf (10:1) 96 h | $(6.0 \pm 0.1) \times 10^6$ | $(7.4 \pm 0.5) \times 10^7$ | $(7.9 \pm 0.5) \times 10^7$ | 1:3.3 | 0.6 | 12.3:1 |
| LS 168 h | $(5.4 \pm 0.1) \times 10^6$ | $(8.5 \pm 0.4) \times 10^7$ | $(9 \pm 0.3) \times 10^7$ | 1:3.9 | | 15.7:1 |
| Co-inf (10:1) 168 h | $(5.1 \pm 0.1) \times 10^6$ | $(8.3 \pm 1.2) \times 10^7$ | $(8.8 \pm 1.2) \times 10^7$ | 1:2.5 | 0.6 | 16.3:1 |

**Table S8. Quantitative analysis of viral load in infected THP-1 macrophages**

Viral load was quantified from infected THP-1 cell culture supernatant and cell lysate at 48, 96, and 168 h.p.i.. Data are presented as mean ± SD from a minimum of three independent experiments.

| Sample | Virus copies/3 mL sup | Virus copies/500 µL lysate | Total Virus gE (EC+IC) | Parasite: Virus Ratio | Fold increase compared to only LS | IC: EC |
|---|---|---|---|---|---|---|
| LS 48 h | 7 (± 0.4) x $10^5$ | 6.7 (± 0.9) x $10^6$ | 7.4 (± 0.9) x $10^6$ | 1:6.1 | | 9.7: 1 |
| Co-inf (10:1) 48 h | 0.7 (± 0.1) x $10^5$ | 1.1 (± 0.1) x $10^6$ | 1.2 (± 0.07) x $10^6$ | 1:6.3 | 1 | 14.8:1 |
| LS 96 h | 6.5 (± 0.1) x $10^5$ | 2.1 (± 0.2) x $10^7$ | 2.1 (± 0.2) x $10^7$ | 1:9.5 | | 31.6:1 |
| Co-inf (10:1) 96 h | 1 (± 0.1) x $10^5$ | 1.4 (± 0.1) x $10^7$ | 1.4 (± 0.1) x $10^7$ | 1:17.7 | 1.7 | 135:1 |
| LS 168 h | 4.8 (± 0.3) x $10^5$ | 2.2 (± 0.2) x $10^7$ | 2.2 (± 0.2) x $10^7$ | 1:3.6 | | 45: 1 |
| Co-inf (10:1) 168 h | 0.9 (± 0.1) x $10^5$ | 1.1 (± 0.01) x $10^7$ | 1.1 (± 0.01) x $10^7$ | 1:2.1 | 0.61 | 130: 1 |
